# Under the name of “Lua”: Revisiting Genetic Heterogeneity and Population Ancestry of Austroasiatic speakers in Northern Thailand Through Genomic Analysis

**DOI:** 10.1101/2024.04.16.589696

**Authors:** Jatupol Kampuansai, Tanapon Seetaraso, Maneesawan Dansawan, Suwapat Sathupak, Wibhu Kutanan, Metawee Srikummool, Angkana Inta

## Abstract

Austroasiatic (AA)-speaking populations in northern Thailand are of significant interest due to their status as indigenous descendants and their location at the crossroads of AA prehistoric distribution across Southern China, the Indian Subcontinent, and Mainland Southeast Asia. However, the complexity of ethnic identification can result in inaccuracies regarding the origin and migration history of these populations. To address this, we conducted a genome-wide SNP analysis on 89 individuals from two Lavue- and three Lwa-endonym populations and combined them with previously published data to elucidate the genetic diversity and clustering of AA groups in northern Thailand. Our findings align with linguistic classifications, revealing distinct genetic structure among the three branches of the Mon-Khmer subfamily within the AA family: Monic, Khmuic, and Palaungic. Although the term “Lua” ethnicity is used confusingly to identify ethnic groups belonging to both Khmuic and Palaungic branches, our genomic data clarifies that the Khmuic-speaking Lua living on the eastern side of the region show genetic differentiation from the Palaungic-speaking Lavue and Lwa populations living on the western side. Within the Palaungic branch, the Dara-ang population stands out as genetically distinct, reflecting remnants of ancient ancestry. The Lavue populations, mainly inhabiting mountainous areas, exhibit a genetic makeup unique to the AA family, with a close genetic relationship to the Karenic subgroup of the ST family. Conversely, the Lwa and Blang populations, residing in lowland river valleys, display genetic signatures resulting from admixture with Tai-Kadai-speaking ethnic groups.

**Author summary:** In the past, many Austroasiatic speakers in northern Thailand concealed their identity due to perceptions of inferiority compared to the majority Tai-Kadai-speaking group in the region. However, attitudes have shifted in the modern era, with a greater appreciation for ethnic diversity and the unique aspects of different valuable cultures. This scenario provides an opportunity for us to use genetic insights to untangle the complexities of ethnic identification among these indigenous inhabitants of Mainland Southeast Asia.

Through our genetic analysis, we aimed to shed light on the ancestry and diversity of the Austroasiatic people in northern Thailand, often collectively referred to as “Lua” or “Lawa”, which is an exonym (a name of the ethnic group created by another group of people) commonly used in prior scientific reports. Our findings clearly indicate genetic distinctions among the Lua, Lavue, and Lwa ethnic groups. The intricate interplay of genetics, cultural heritage, and historical influences has shaped these ethnic communities. Our study underscores the importance of accurate ethnic classifications, emphasizing the use of self-identified endonyms, names created and used by the ethnic group themselves. This approach respects the Austroasiatic communities in northern Thailand and acknowledges their significant contributions to advancing our understanding of genetic anthropology.

## Introduction

The Austroasiatic (AA)-speaking people are ethnolinguistic group in Southeast Asia and parts of South Asia, with a diverse history spanning millennia, encompassing various cultures, languages, and societies. The AA language family stands out as a primary linguistic group in Southeast Asia, alongside Tai-Kadai (TK), Sino-Tibetan (ST), Austronesian (AN), and Hmong-Mien (HM) languages. The ancestors of AA-speaking populations were likely Neolithic rice farmers residing in southern China around 7,000 years ago [1]. This population embarked on multiple migration waves into Southeast Asia, a process that began as early as the Neolithic era [2,3]. This migration was likely driven by a combination of factors, including population growth, environmental shifts, and the expansion of agricultural practices [4]. The development of settled agricultural societies, facilitated by the spread of agriculture, played a pivotal role in the emergence of complex civilizations in Southeast Asia. One notable example is the Dvaravati Mon civilizations that flourished around the 7th century AD in Thailand’s Chao Phraya valley, a region renowned for its rice production [5].

The northern part of Thailand holds strategic importance as it serves as a vital gateway connecting Thailand to neighboring countries such as Myanmar, Laos, and China. This area has been a historical crossroads for the southward migration of AA people from East to Southeast Asia since ancient times. In northern Thailand, AA-speaking populations are recognized as indigenous groups descended from prehistoric populations [6]. All of them are linguistically classified within the Mon-Khmer subfamily of the AA family, further divided into three branches: Monic, Khmuic, and Palaungic [7]. The Monic branch comprises only the Mon ethnicity, while the majority of present-day AA speakers in northern Thailand belong to the Palaungic and Khmuic branches, categorized under the Northern Mon-Khmer subfamily, with estimated populations of 20,483 and 58,405 people, respectively [8].

Among the identification of AA-speaking ethnicities residing in northern Thailand, the term “Lua” is the most confusing in various research works because it is used to refer to certain ethnic groups in the Khmuic branch living in the eastern part of the northern region near the border with Laos. Simultaneously, some researchers use it to describe the Palaungic-speaking community in the highland areas of Chiang Mai and Mae Hong Son provinces, located in the western part of the northern Thailand near the border with Myanmar (Fig. 1 and Table 1). In fact, each of these ethnicities has their own endonyms (names created and used by the ethnic group itself) and differs significantly in terms of ancestry and historical background. The confusion surrounding the term “Lua” arises from its similarity to closely related ethnonyms, such as Lua, Lawa, Lavue, and Lavua [9]. Since they all speak languages belonging to the AA family, outsiders often overlook these differences and assume they are the same ethnic group. For example, “Lua” or “Lawa” is an exonym (a name of the ethnic group created by another group of people) commonly used by Northern Thai, Central Thai, and Christians to refer to Palaungic-speaking people, but it is not an endonym used by these people to identify themselves. Furthermore, in some historical contexts, the term “Lua” does not specifically denote any ethnic group but rather serves as a derogatory term used by rulers to distinguish indigenous inhabitants residing outside urban areas [10].

**Table 1.** The information pertains to the AA-speaking population residing in northern Thailand, for whom genome-wide autosomal data is accessible.

| Code | Ethnic name |  | Location<br>(village, district,<br>province) | Language<br>branch | No. of<br>samples | Reference |
| --- | --- | --- | --- | --- | --- | --- |
|  | endonyms | exonyms |  |  |  |  |
| Lwa1 | Lwa*, Lua | Lua, Lawa | Khun Khong, Hang<br>Dong, Chiang Mai | Palaungic | 20 | This study |
| Lwa2 | Lwa*, Lua | Lua, Lawa | U-Meng, San Pa<br>Tong, Chiang Mai | Palaungic | 15 | This study |
| Lwa3 | Lwa*, Lua | Lua, Lawa, Pareak | Nong Khiew, Chiang<br>Dao, Chiang Mai | Palaungic | 14 | This study |
| Lavue1 | Lavue, Laveue | Lua, Lawa | Hao, Mae Cham,<br>Chiang Mai | Palaungic | 25 | This study |
| Lavue2 | Lavue, Laveue | Lua, Lawa | Meuad Long, Mae<br>Cham, Chiang Mai | Palaungic | 15 | This study |
| Lavue3 | Lavue, Laveue | Lua, Lawa<br>(Western) | Dong, Mae La Noi,<br>Mae Hong Son | Palaungic | 10 | Changmai et al.,<br>2022 |
| Lavue4 | Lavue, Laveue | Lua, Lawa<br>(Western) | Pa Pae, Mae Sariang,<br>Mae Hong Son | Palaungic | 9 | Kutanan et al.,<br>2021 |
| Lavue5 | Lavue, Laveue | Lua, Lawa<br>(Eastern) | Bo Luang, Hod,<br>Chiang Mai | Palaungic | 10 | Kutanan et al.,<br>2021 |
| Blang | Blang | Lua, Blang, Tai<br>Doi, Sam Tao | Lua Pattana, Mae<br>Chan, Chiang Rai | Palaungic | 9 | Kutanan et al.,<br>2021 |
| Dara-ang | Dara-ang | Palaung | Nor Lae, Chiang Dao,<br>Chiang Mai | Palaungic | 9 | Kutanan et al.,<br>2021 |
| Lua1 | Lua | Lua, Htin | Huay Lom, Bo Kluea,<br>Nan | Khmuic | 13 | Kampuansai et<br>al., 2023 |
| Lua2 | Lua | Lua, Htin | Pa Kam, Bo Kluea,<br>Nan | Khmuic | 14 | Kampuansai et<br>al., 2023 |
| Lua3 | Lua | Lua, Htin, Mal | Ta Luang, Pua, Nan | Khmuic | 10 | Lipson et al.,<br>2018 |
| Lua4 | Lua | Lua, Htin, Pray | Huay Kaew, Chiang<br>Klang, Nan | Khmuic | 11 | Kutanan et al.,<br>2021 |
| Khamu1 | Kmhmu, Khamu | Khamu, Khammu | Pang Sa, Tha Wang<br>Pha, Nan | Khmuic | 17 | Kampuansai et<br>al., 2023 |
| Khamu2 | Kmhmu, Khamu | Khamu, Khammu | Chon Dan, Song<br>Khwa, Nan | Khmuic | 16 | Kampuansai et<br>al., 2023 |
| Khamu3 | Kmhmu, Khamu | Khamu, Khammu | Huay Puk, Mueng,<br>Nan | Khmuic | 21 | Kampuansai et<br>al., 2023 |
| Khamu4 | Kmhmu, Khamu | Khamu, Khammu | Huay Sataeng, Thung<br>Chang, Nan | Khmuic | 10 | Kutanan et al.,<br>2021 |
| Mlabri | Mlabri | Mlabri | Huay Yuak, Wiang<br>Sa, Nan | Khmuic | 9 | Lipson et al.,<br>2018 |
| Mon_NT | Mon | Mon | Nong Du, Pa Sang,<br>Lamphun | Monic | 9 | Changmai et al.,<br>2022 |

\* The present-day villagers pronounce their endonym as “Lua”. However, to distinguish them from the “Lua” ethnicity in Nan province, we choose to use “Lwa”, which is their historic ethnonym.

**Fig. 1.**
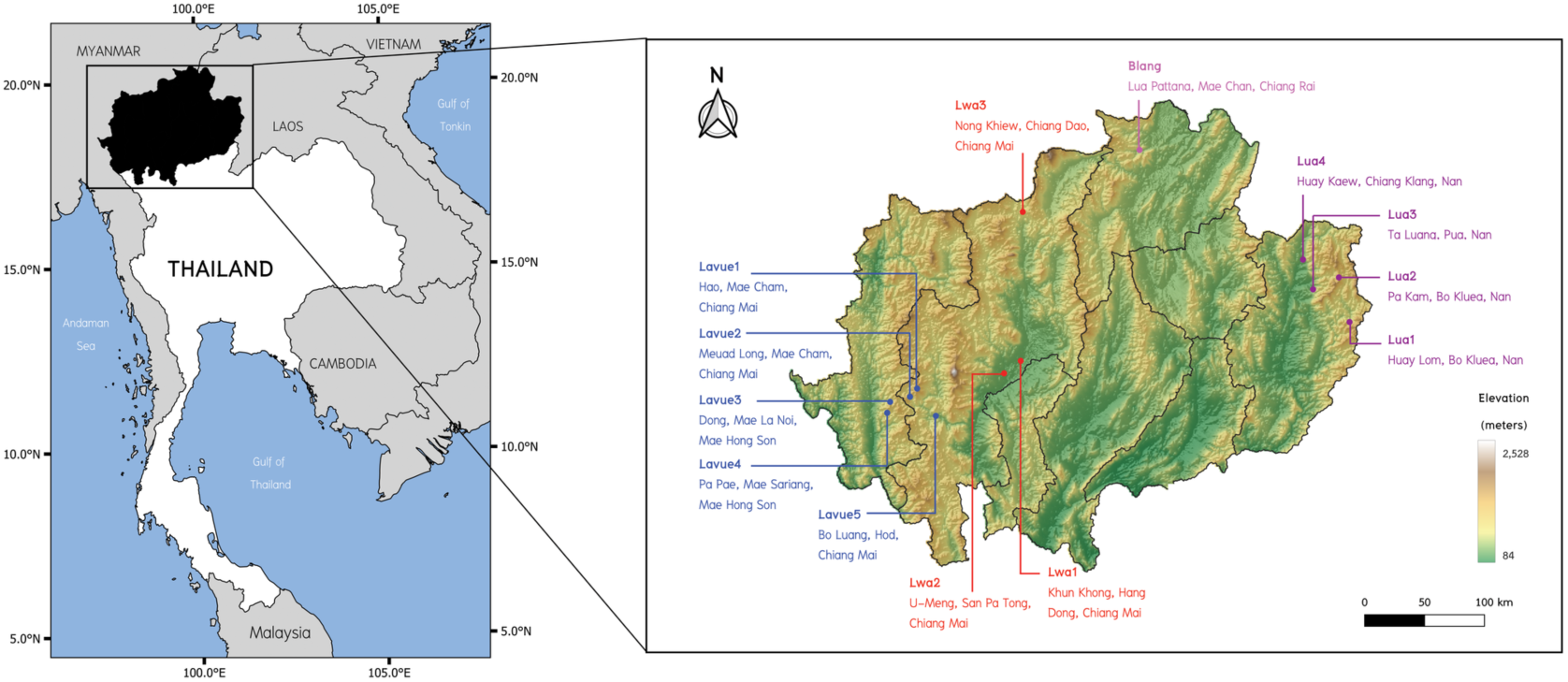
Geographical distribution of AA samples commonly referred to as the Lua collective name. The colored text on the map of northern Thailand represents the Lua or related ethnonym. Background map was created using QGIS 3.6.0 (http://www.qgis.org/).

This misidentification of ethnicities has implications for genetic anthropology studies, as some research may incorrectly consider these distinct groups as the same ethnicity and analyze data without recognizing their differences. This issue is particularly significant because the AA people, who are the original inhabitants of Mainland Southeast Asia, represent a population that could be highly relevant to the extensive ongoing exploration in ancient DNA studies. While some previous studies have attempted to clarify the genetic classification of AA speakers in northern Thailand [11,12], the limited number of samples in the Palaungic branch, which is the most diverse and complex within this linguistic family, has hindered the revelation of their genetic ancestry and history. Therefore, we have generated new genome-wide data for AA-speaking populations in northern Thailand, specifically targeting all ethnic endonyms within the Palaungic branch, aiming to elucidate the intricacies of subgroup classification within the AA linguistic family.

## Results

### Population structure

We generated genome-wide SNP data from 89 Palaungic-speaking individuals, belonging to the Lavue and Lwa ethnonyms, residing in northern Thailand (Table 1, Fig. 1). Using a set of 219,244 SNP positions, we conducted Principal Component Analysis (PCA) to explore the genetic structure and relationships among a total of 2,355 individuals, including our samples and various Asian populations (Fig. 2). The PCA analysis divided the population into subgroups based on geographical regions. PC1 distinguished East Asians (on the left side) from South Asians (on the right side), while PC2 separated Southeast Asians (at the top) from Northeast Asians (at the bottom). Additionally, when considering subgroups by linguistic families, populations speaking the same language tended to cluster together, except for the AA family, which was dispersed across the map. In our newly generated data, we observed that the Palaungic language group showed genetic proximity to the AA, TK, and ST families, with substantial overlap in their positions. Within the Palaungic branch, the Lavue, Lwa, and Blang samples were closely positioned with overlaps, while the Dara-ang people exhibited distinct positions.

**Fig. 2.**
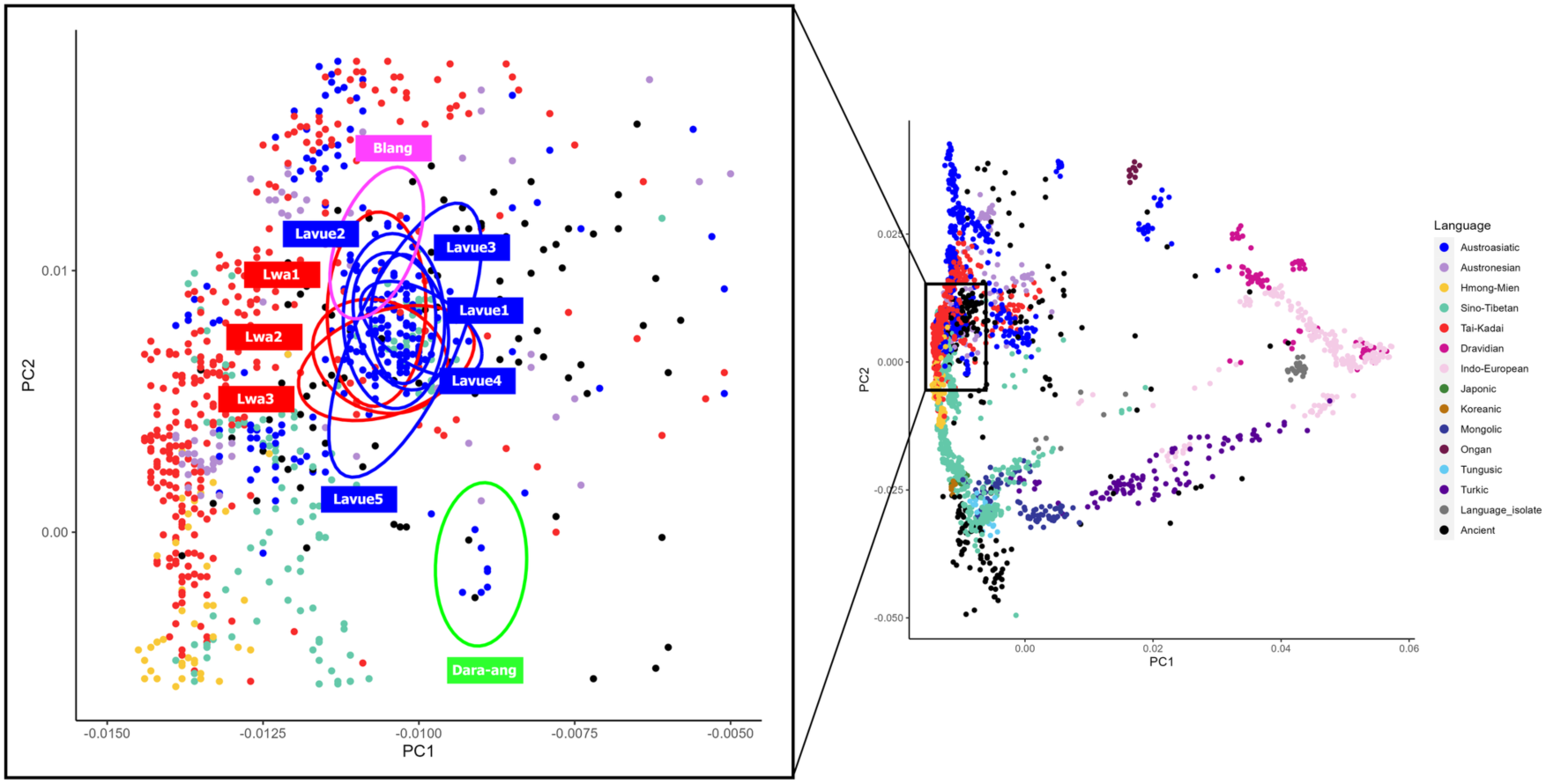
PCA plot for the genome-wide SNP data of individuals from East and South Asia. Each individual is colored by linguistic family according to the key in the right panel. The plot focusing on the Palaungic-speaking populations is zoomed in on the left.

We proceeded to examine the genetic components using ADMIXTURE program version 1.3.0 [13], dividing the gene pool into groups (K) ranging from 2 to 12, with each grouping repeated 100 times. The analysis indicated that the lowest cross-validation value occurred at K=9, suggesting that 9 groups represent the most suitable number of ancestral components (S1 Fig.).

At K=9 (Fig. 3), when considering linguistic family groups, it’s evident that the AA family exhibits a specific genetic component in purple, while TK and AN display a major yellow component, and HM shows the highest proportion of blue-green components. The ST family is characterized by a prominent blue component, whereas red and pink components are primarily associated with Dravidian and Indo-European language families. Moreover, some populations demonstrate unique proportions; for instance, the Mbuti people showcase a dominant green component, the Mani people have a major gray component, and the Mlabri people exhibit a major orange component.

**Fig. 3.**
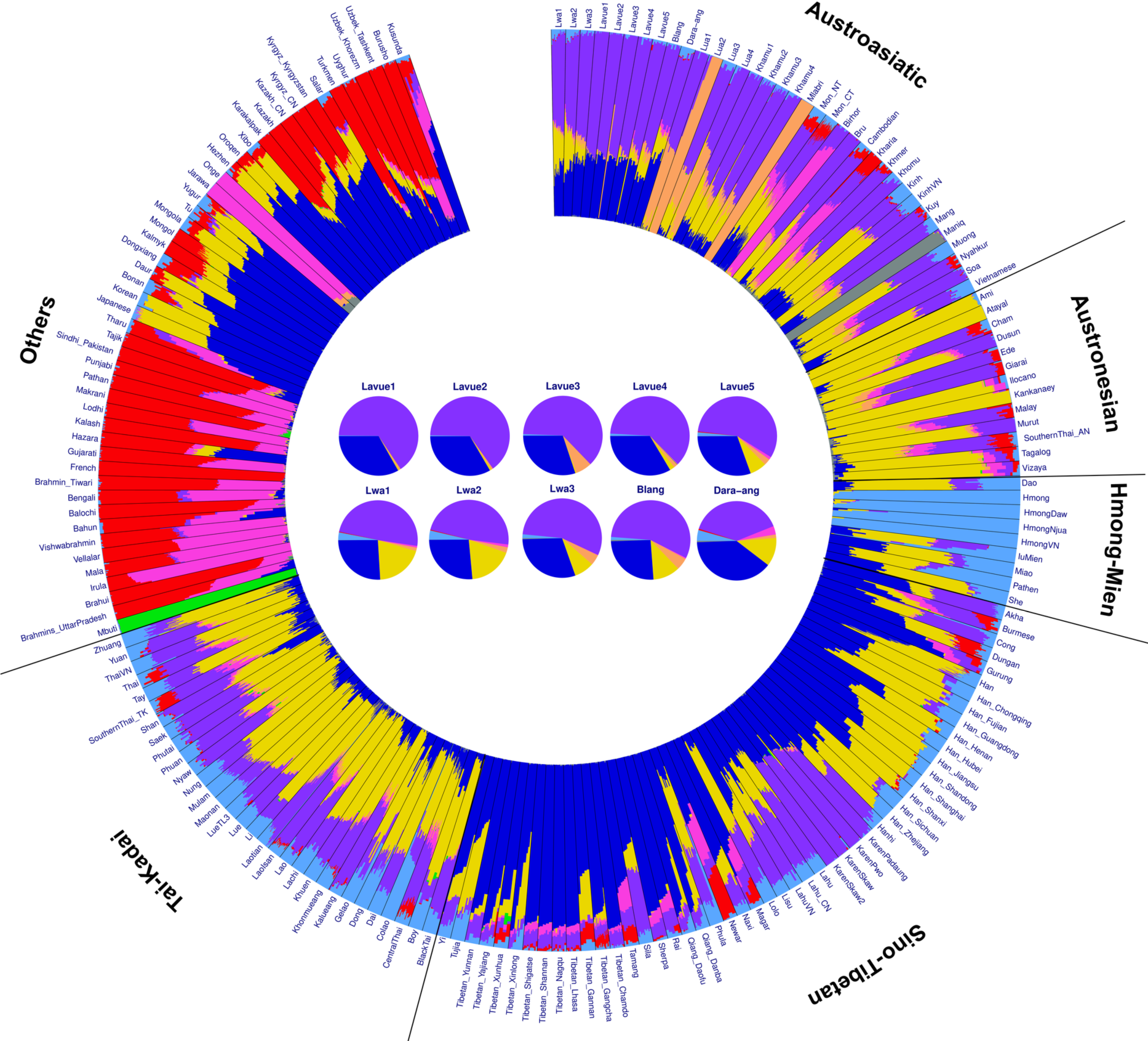
The ADMIXTURE results of modern populations in from East and South Asia. The plot delineated into K=9 groups of ancestral components. Each individual is depicted by a bar segmented into K colored sections, representing their estimated ancestry components. Populations are demarcated by black lines, and their linguistic family is labeled outside the diagram.

Regarding the Palaungic branch (Fig. 3), our findings reveal that the Lwa, Blang, and Dara-ang populations exhibit three main genetic components (purple, blue, and yellow), while the Lavue population has two primary genetic components (purple and blue), resembling the genetic components of the ST family, which includes the Karen, Lahu, and Lisu ethnic groups. Even as K increased to 12 (S2 Fig.), the Lavue and those speaking ST languages continued to share similar proportions of genetic components. Additionally, upon comparison with ancient DNA, the pink component found in some populations of the AA family was observed in several ancient samples spanning from the Neolithic to Bronze age, albeit in varying proportions (S2 Fig.).

### Genetic Relationships Among Populations

The analysis of genetic relationships among populations involved calculating *f3* and *f4*-statistics. The *f3*-statistics analysis, represented as *f3*(X, Y; Mbuti), quantified the shared drift between two populations, X and Y, since their separation from an African Mbuti outgroup. A higher *f3* value indicates a closer genetic relatedness between populations. Among the AA populations in northern Thailand, specifically those belonging to the Palaungic branch (including Lavue, Lwa, Blang, and Dara-ang), they formed a distinct cluster separate from the Monic (Mon_NT) and Khmuic (Lua, Khamu, Mlabri) branches (Fig. 4 and S3 Table). Notably, the Lavue populations exhibited allele sharing with the KarenPaduang, while the Lwa showed genetic relatedness with certain TK-speaking (Shan and KhonMueang) and ST-speaking (Akha, Lisu, Lahu) groups.

**Fig. 4.**
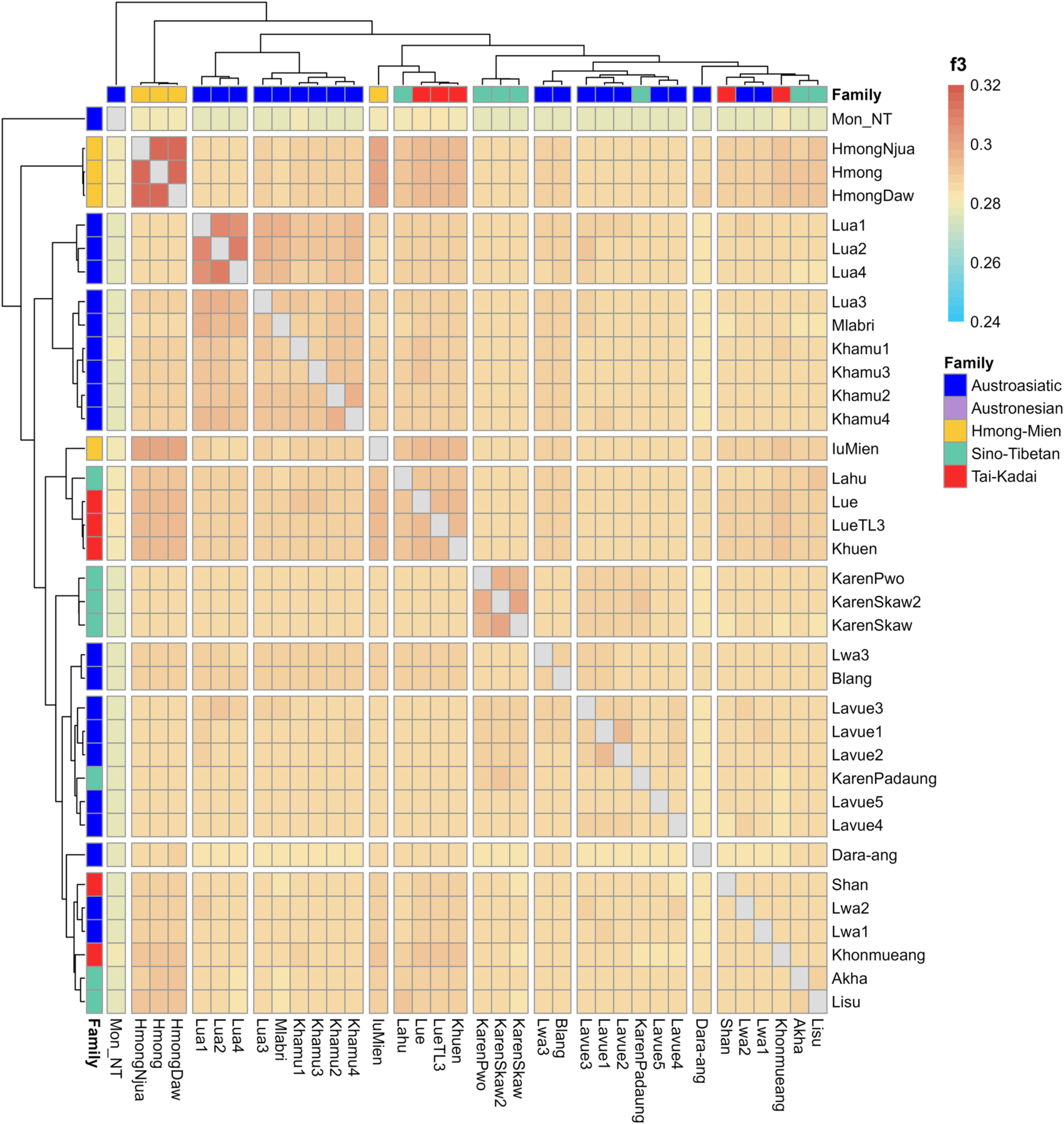
The heatmap displays allele sharing profiles of populations in northern Thailand, as determined by *f3*-statistics. The colored bar located at the top-right corner indicates the statistical values and linguistic family of each ethnic group.

When applying the *f3*(X, Y; Mbuti) form to modern populations in Asia (represented by X) and the AA-speaking group (represented by Y), the analysis revealed close genetic relatedness within the AA language group, with some exceptions like the Maniq people and certain populations from India (Birhor and Kharia) (S3 Fig.). The clustering pattern of the AA-speaking people in northern Thailand aligned with their endonyms as Lua, Lavue, and Lwa. Upon comparison with ancient DNA, the Dara-ang people were genetically distinct from their Palaungic-speaking relatives (S4 Fig.). Interestingly, all ethnic groups belonging to the Palaungic branch in northern Thailand and some AA-speaking populations from Vietnam (such as Muong, Kihn, Vietnamese) showed the closest genetic affinity with a Neolithic sample from Wuzhuang Guoliang in China, recognized as representative of Yellow River Farmers [14] (S4 Fig.).

When examining population relationships using *f4*-statistics in the format *f4* (W, X; Y, Mbuti), where W represents a specific Palaungic-speaking Dara-ang ethnic group, X denotes other populations within the Palaungic branch, and Y encompasses various ethnic groups from the AA, TK, ST, AN, and HM families (S4 Table), the proximity of points to the compared ethnic groups (as depicted on the left side of Fig. 5) indicates close genetic relationships. Generally, populations within the Palaungic branch demonstrated a close genetic affinity with subgroups of the AA family, particularly the Khmuic (including Khamu, Mlabri, and Lua) and Katuic (Kui, Bru, and So) branches. They also exhibited genetic relatedness with the Karenic branch of the ST family and certain AN-speaking populations (such as Ede and Giarai) in Vietnam.

**Fig. 5.**
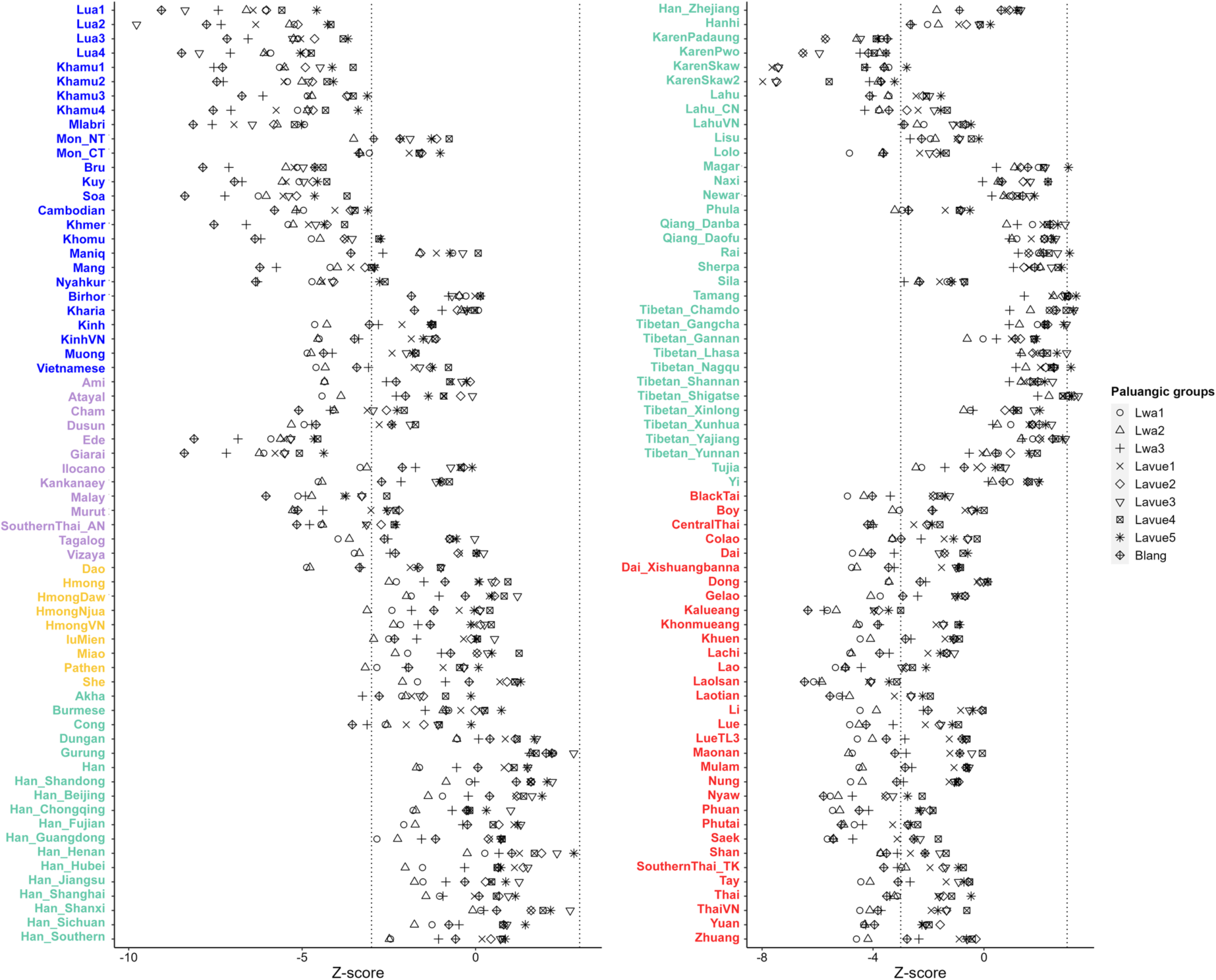
The *f4*-statistics compare Palaungic-speaking populations with ethnic groups from various language families. Z-scores are computed for *f4*(W, X; Y, Mbuti), where W represents a selected AA-speaking Dara-ang ethnic group, X represents other populations from the Palaungic branch, and Y denotes a population not in the Palaungic branch, with their ethnic names labeled on the left side. Different symbols are used to represent different populations for X. The ethnic names are labeled in colors to show different linguistic family. The vertical black dashed lines denote +3/-3.

In comparison to the TK family, we observed that the Lwa and Blang populations tended to have a closer relationship with TK-speaking populations than the Lavue population (Fig. 5). Additionally, when comparing the genetic data of various modern AA-speaking populations with ancient DNA from Southeast Asia, all populations of the Khmuic branch showed relatedness with samples from TamPaling in Laos and TamYappaNhae in Thailand. Interestingly, Lavue3 is the only one population in the Palaungic branch who displayed significant genetic affinity with Iron Age ancient DNA from the TamYappaNhae archaeological site in Mae Hong Son province of Thailand (S5 Fig. and S5 Table).

### Genetic ancestry of populations

The genome-wide data of AA speakers in Northern Thailand and various Asian ancestral populations, including those outside the region, were analyzed using the TreeMix version 1.12 program [15] to construct a maximum-likelihood tree. The results illustrate genetic distinctions among population groups belonging to the Khmuic and Palaungic branches in northern Thailand (Fig. 6). Specifically, within the Palaungic branch, Lavue1-4 form a cluster along with KarenSkaw and KarenPwo, while Lavue5 clusters with KarenPadaung. The Lwa3 clusters with the Blang ethnic group, whereas the other Lwa populations exhibit diversity within the tree. When we incorporate migration events into TreeMix analyses, we observe a connection between the clade of Khmuic-speaking people in Thailand and a clade comprising some ethnic groups from Vietnam (Vietnamese and Khomu) (Fig. 6).

**Fig. 6.**
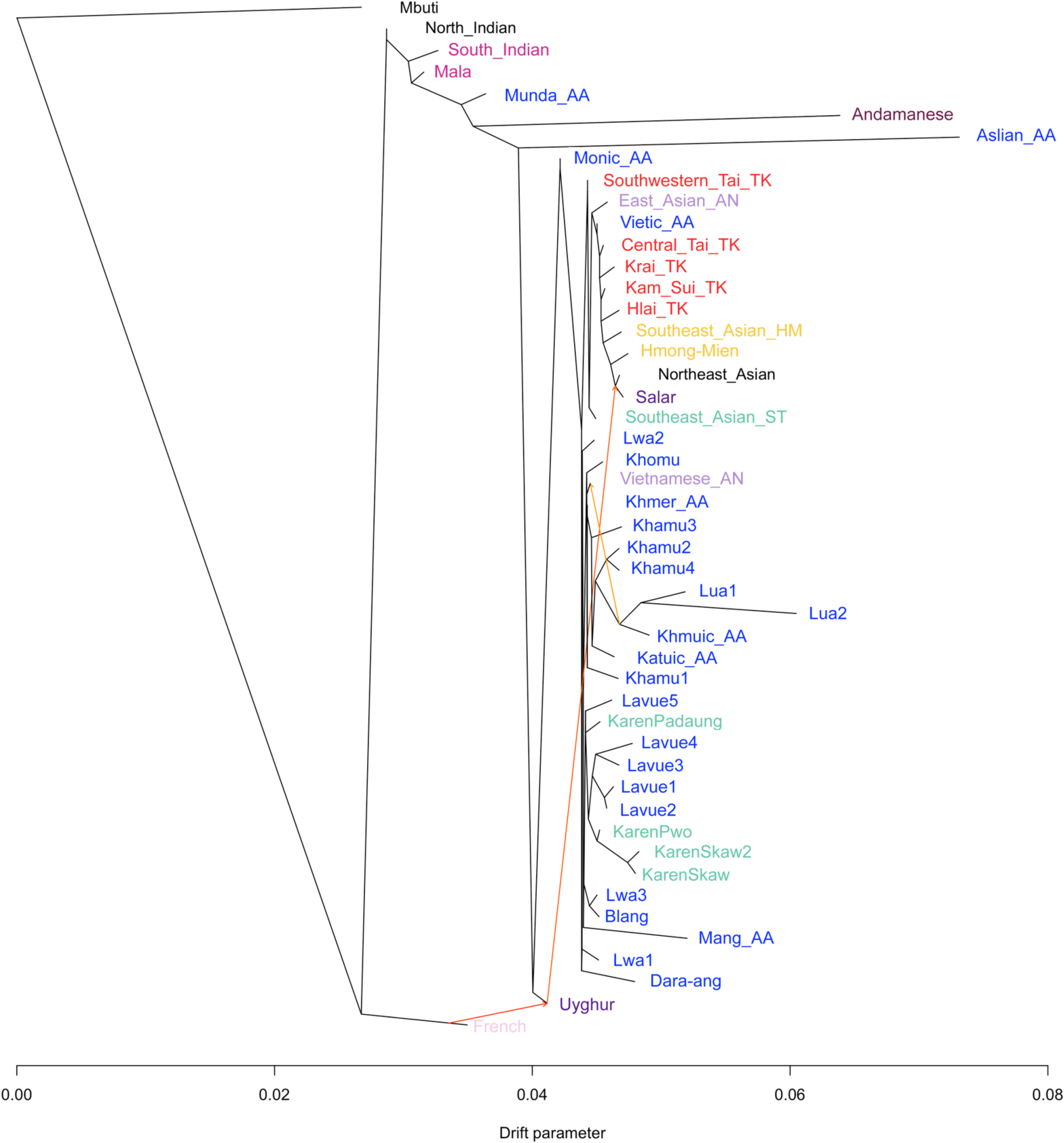
The TreeMix diagram with three migration events of the AA-speaking ethnic group in northern Thailand and other modern East Asia. Populations are labeled with different colors based on their language family.

We delved deeper into ancestry by initially constructing backbone admixture graphs using representatives from the AA, TK, AN, HM, and ST language families, namely the Cambodian, Dai, Atayal, Miao, and Naxi, respectively, with the Mbuti serving as the outgroup. Then, we incorporated AA speakers from northern Thailand into the analysis and found the best-fitting graph (Fig. 7b). Upon isolating the outgroup, the AA-related ancestry separated from the North Indian and emerged as the Dara-ang, followed by the Blang. The Lwa presented as admixed populations, incorporating AA-related and Dai-related ancestries in proportions of 38% and 62%, respectively. Subsequently, we delved further to determine the ancestry of the Lavue and Karen populations, which showed genetic closeness. We incorporated Lavue, Karen, and Dara-ang into the graph, grouping them based on TreeMix clustering: Lavue (Lavue1-4), Lavue5, KarenPaduang, Karen (KarenSkaw and Karen Pwo), and Dara-ang. We observed shared ancestry between the Lavue and Karen populations. This ancestral lineage then diverged to form the Dara-ang group, which exhibited a greater influence from North Indian-related ancestry. The Lavue populations emerged as a result of a mixture between Lavue-related and Karen-related ancestries, with a ratio of 70% and 30%, respectively (Fig. 7c).

**Fig. 7.**
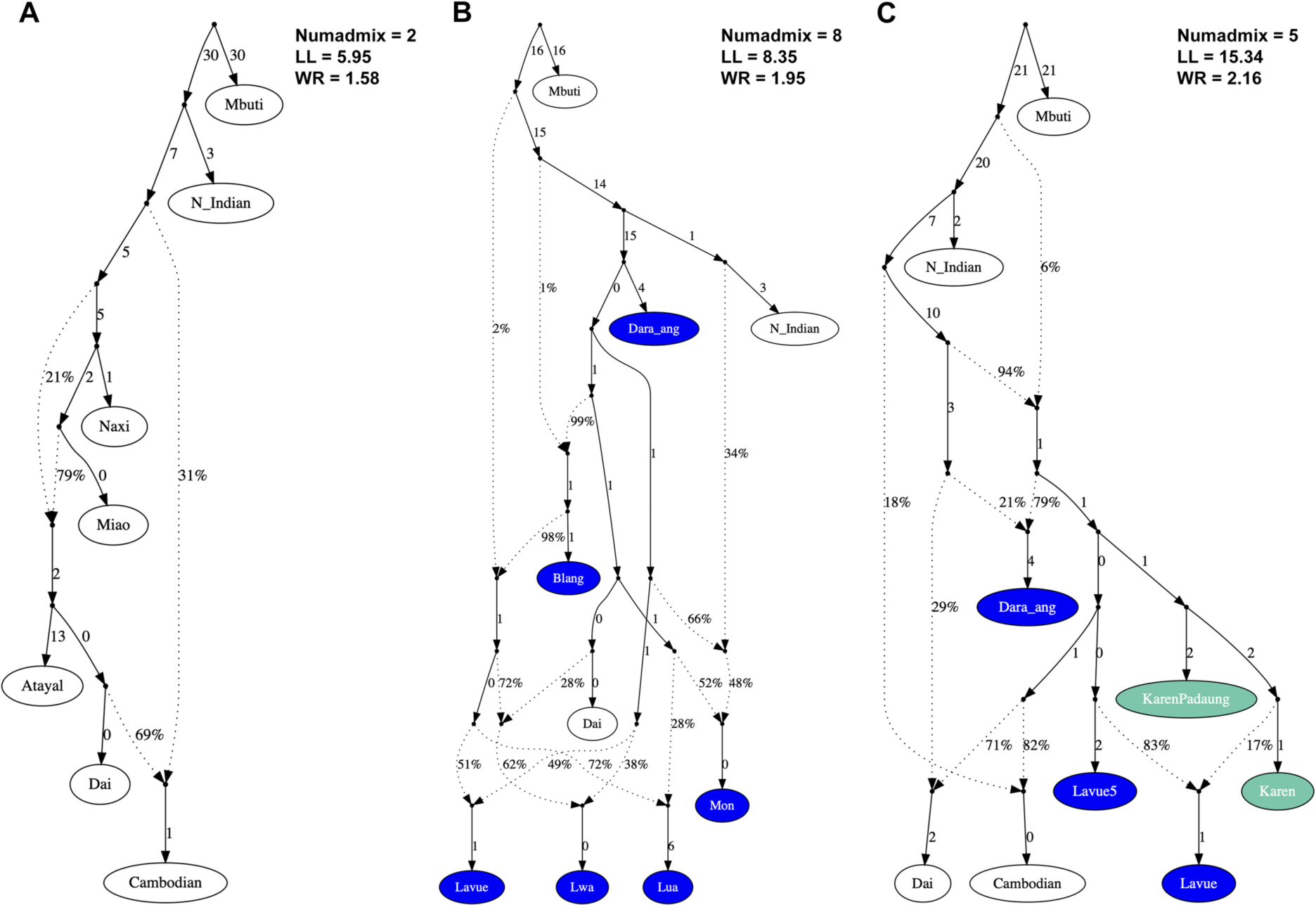
Admixture graphs of AA populations. (A) representatives of different language families including the Atayal, Dai, Cambodian, Miao, and Naxi for the AN, TK, AA, HM, and ST language families, respectively; (B) AA-speaking groups of northern Thailand; (C) Palaungic- and Karenic-speaking groups. White nodes denote backbone populations. Dashed arrows represent admixture edges, while solid arrows indicate drift edges reported in units of FST × 1,000.

## Discussion

In the past, many AA-speaking individuals in northern Thailand felt embarrassed about their identity and often tried to conceal it by assimilating into the majority Northern Thai population [10]. This situation created confusion for historians and scientists studying these communities. However, with the growing acceptance of ethnic diversity in the modern world, this trend is changing. While these individuals consider themselves Thai, they also take pride in their unique culture and language. This changing presents an opportunity to clarify and correct the AA ethnic identity, and we encourage the scientific community to use their self-identified ethnonyms, acknowledging their valuable contributions to the field of genetic anthropology.

The populations of AA-speaking language in northern Thailand are linguistically linked with a structure within the Mon-Khmer subfamily [7], suggesting a shared ancestry that has persisted over time, evident in their close relationships and similar genetic components as observed in our study (Fig. 2). However, upon further examination of their genetic sub-structure, differences emerge among these AA populations, which we will delve into based on their specific language branches.

### Monic branch

The Mon people are historically recognized as one of the earliest settlers in Mainland Southeast Asia, with their civilization centered in Central Thailand and Southern Myanmar dating back over thousand years. They are renowned for their sophisticated art, architecture, Buddhism, and literary works, which have significantly influenced the cultural development of the Thai and Burmese people. The Mon established the renowned Dvaravati kingdom, spanning from Southern Burma (Myanmar) to Central Thailand between the 3rd and 10th centuries AD. By the 8th century AD, their civilization extended to the Haripunchaya region in present-day Lamphun province, Northern Thailand [8]. However, due to displacement and assimilation by the TK-speaking groups over centuries, only a small number of individuals still identify as Mon in contemporary times.

Although the Mon and the local inhabitants referred to as “Lua” have been consistently mentioned together throughout the history of northern Thailand [16,17], they have generally been recognized as distinct ethnic groups. The differences in their languages and genetic structures clearly distinguish the Mon of northern Thailand from other AA-speaking ethnicities. The remaining Mon population in northern Thailand still maintains similarities with the Mon in the central region, evident through their southern Mon-Khmer language, cultural customs, Buddhist practices, and the genetic relatedness we have recently observed (S3 Fig). This genetic distinction between the Mon and other AA-speaking populations also reflects in the influence of South Asian admixture found among the Mon of northern Thailand, which occurred approximately 600 years ago [18].

### Khmuic branch

The AA-speaking population in northern Thailand, belonging to the northern Mon-Khmer subfamily, can be broadly categorized into two primary branches: the Khmuic branch, concentrated mainly along the Thailand-Laos border with a notable presence in Nan province, and the Palaungic branch, primarily situated in Chiang Mai and Mae Hong Son provinces in the western part of the region (Fig. 1). Some ethnicities from these two branches are collectively referred to as the “Lua.”, and it remains an open question whether they have any kin relations [19]. However, these groups differ significantly in terms of language, belonging to distinct branches of the AA family, as well as in cultural practices and post-marital residence patterns. Traditionally, Lavue communities in Chiang Mai and Mae Hong Son follow a patrilocal residence system, where the bride moves to live with the husband’s family after marriage, while Lua communities in Nan practice a matrilocal residence system, tracing their lineage through the female lineage [8].

Our genetic analysis compared the Khmuic and Palaungic branches, revealing distinct differences among the Lua, Lwa, and Lavue groups. The Lua people in Nan province share genetic heritage with local Khmuic-speaking populations like the Htin, Khamu, and Mlabri. Our study also supports the genetic affinity between Khmuic-speaking communities and the Katuic, another AA branch in northeastern Thailand [20]. Comparisons with other modern populations and ancient DNA in Southeast Asia show relatedness between the people of Khmuic branch and AA-speaking Mang and Khomu ethnicities in Vietnam (S4 and S6 Fig.), as well as connections to ancient DNA from the TamPaLing Bronze-age site in northern Laos and the YapPaNhae Iron-age site in northern Thailand (S5 Fig.). While direct historical connections are unclear, the distribution of Khmuic-related ancestry across modern and ancient populations in Thailand, Laos, and Vietnam supports the hypothesis that they share ancient ancestry [12,21,22]. The evidence of a genetic connection observed between the Khmuic population residing in the eastern part of Northern Thailand and ethnic communities in Laos and Vietnam was absent in the Palaungic-speaking groups in the western part of the region. This difference could contribute to the genetic differentiation between the people of the Khmuic and Palaungic branches.

In line with previous findings [12], we confirm that the Lua people in Nan province are the same ethnic group previously identified as the Htin in earlier genetic studies [2,23–26]. The confusion regarding calling the Lua people as the Htin ethnicity stems from differing perspectives. Outsiders, government officers, or other lowland communities typically refer to the indigenous people in this area as Htin (meaning “local people” in literal term). However, the Lua people identify themselves as Lua, and many of them find being called Htin derogatory. Although the term Lua is preferred out of respect for the community, the term Htin is still officially recognized as a hill tribe by the Thai government [27]. It’s also important to note that some works use alternative names for the Lua (Htin) in Nan province, such as Mal and Pray (Prai), which linguistically denote two subgroups of the Lua. While they are distinguished by their ceremonial practices and suggested to have separated lineage-wise a few hundred years ago [28], there is currently no genetic evidence available to differentiate these two Lua subgroups.

### Palaungic branch

The majority of Palaungic speakers in northern Thailand are concentrated in Chiang Rai, Chiang Mai, and Mae Hong Son provinces, residing in mountainous areas near the Myanmar border. Our analysis using PCA and *f3*-statistics (Fig. 2 and 4) revealed close genetic affinities among most Palaungic-speaking communities, except for the Dara-ang people. Examination of genetic components showed that all studied Palaungic-speaking populations have significant genetic contributions from two main sources: the purple component, commonly found in AA-speaking groups, and the blue component, prevalent in ST-speaking groups, particularly the Tibeto-Burman subfamily (Fig. 3 and S2 Fig.).

#### Dara-ang

The Dara-ang is the endonym of the people who are usually known as Palaung, a Burmese language exonym. In China, they are known as De’ang and live in the southwestern part of Yunnan province. The Dara-ang are believed to be the first settlers on the upper bank of the Salween River before the arrival of any other Palaungic-speaking people [8]. There are now more than 250,000 people in Myanmar, but only about 1,937 persons in Thailand, mostly immigrants who moved in 1983 AD and settled in Chiang Mai province, next to the country’s border. Although there are many subgroups of the Dara-ang, such as Pale (or Silver), Shwe (or Golden), Rumai, Riang, all those in Thailand belong to the Pale Dara-ang [8].

Our genome-wide analysis has revealed that the Dara-ang group stands out as the most distinct among the Palaungic relatives (Fig. 2 and 4). The ADMIXTURE diagram reveals that the Dara-ang possess unique pink genetic elements, a feature uncommon in other Palaungic-speaking ethnicities (Fig. 3). When compared with Asian populations, this pink genetic component is akin to that found in the Onge, Jarawa, and Birhor tribes, indigenous hunter-gatherer communities inhabiting the Andaman Islands and the Indian subcontinent. It’s plausible that some genetic lineages trace back to the earliest Southeast Asian migrants who journeyed through the Indian subcontinent, persisting within the Dara-ang gene pool. However, our examination using *f4*-statistics and ancient DNA comparisons (S5 Fig.) did not establish a close link between the Dara-ang and ancient DNA sources across Southeast Asia. This might be attributed to the limited availability of ancient DNA data in this region. Extending the comparison to include more ancient DNA data from Southeast Asia and the Indian subcontinent could provide clarity regarding the origins of this particular genetic lineage observed in the Dara-ang population.

#### Lavue

Since the Lavue and Lwa lack a written language, there are no documented records detailing their settlements and migrations. However, oral traditions recount a narrative suggesting that the Lavue and Lwa were originally part of the same ethnic group that once inhabited a city in the Ping River plain, situated in the modern-day Chiang Mai-Lamphun basin of northern Thailand. They then were overcome by the incoming Tai-Kadai (TK) speaking migrants, which resulted in the Lavue migrating to the mountains while the Lwa remained under TK governance. Historical documents from later periods refer to these indigenous peoples by various names such as Lua, Lawa, Milakhu, Tamilla, and La [17].

In our research, we primarily focused on using the self-identification endonyms to categorize the Palaungic branch. Among those previously identified as part of the “Lua” ethnicity, we discovered two major sub-groups: the Lavue residing in mountainous areas and the Lwa in lowland river plains. Through analysis of genome-wide autosomal markers, we observed distinct genetic structures and ancestries between these two sub-groups. The Lavue maintained a genetic structure characteristic of the AA family (represented by purple and blue colors), with minimal genetic admixture with other ethnic groups (Fig. 3 and S2 Fig.). While linguistic evidence indicates dialectical variations within the Lavue’s language (described as the Western Lawa in [10]) with up to eight dialects, our genetic findings reveal shared ancestry and similar genetic components within this group.

We observed an intriguing genetic affinity between the Lavue and the Karen ethnic group of the ST family (Fig. 6). Despite cultural and linguistic differences between the Lavue and Karen communities, they have developed close proximity over time. In the past 150 years, Sgaw Karen individuals from Myanmar have migrated into the Lavue mountains in Thailand, then coexisting with the Lavue population [10]. While intermarriage with non-Lavue individuals has traditionally been rare due to the Lavue community’s adherence to their habitat and culture, such limitations have gradually diminished in contemporary times. Evidence of intermarriages between Lavue individuals and spouses from different ethnic groups, such as Sgaw Karen and Northern Thai, supports this trend [9].

#### Lwa

The Lwa group represents the Palaungic-speaking indigenous population residing in the lowland river plains of northern Thailand. Linguistically, most Lwa individuals are clustered with the Eastern Lawa and have been noted to understand the Lavue’s language at only about 50% comprehension [10]. Our results have also revealed that the genetic structure of the Lwa group in Chiang Mai province differs from that of the Lavue group (Fig. 4). This difference is attributed to genetic admixture, evidenced by the presence of genetic composition from the TK family, which is rare in the Lavue group (Fig. 3 and S2 Fig.). The historical relationship between the Lwa people and TK-speaking groups is well-documented in various historical documents, which describe the Lwa as the original inhabitants of the Chiang Mai-Lamphun basin before the incoming migration of the TK people around the 13th century [6]. The Lwa who still exist in the river plains cannot maintain their original AA culture and have been acculturated and assimilated by the TK community living around them [8]. While some Lwa communities in different districts near Chiang Mai city still acknowledge themselves as the Lwa, they have largely adopted the culture of the TK community and may eventually lose their identity in the future.

It appears that the genetic distinction between the Lavue and Lwa aligns well with their classification based on spoken language, as the Western Lawa [ISO code: lcp] and Eastern Lawa [ISO code: lwl], respectively [7]. However, there is an exception with the Lavue5 from Bo Luang village, Chiang Mai province, which does not fit this categorization. The residents of this village identify themselves with the Lavue endonym, similar to those in the mountainous areas, but they are linguistically classified as Eastern Lawa alongside the Lwa in the lowland river plain. Our analysis also indicates that the genetic structure of Lavue5 resembles that of the Lwa, showing a high degree of admixture with the TK gene pool. This suggests that, within the Palaungic branch, self-identification endonyms and linguistic classifications do not solely reflect genetic relatedness.

Interestingly, we compared our genetic results with animism practices and revealed a correlation between genetic clustering patterns of Palaungic speakers and differences in spiritual worship practices within their communities. Lavue populations with minimal or no evidence of TK admixture (Lavue1-4) (Fig. 3) primarily engage in the spirit ceremony centered around the “Sao Sa Kang” (the village’s sacred pillar), a landmark of animism. On the other hand, populations showing evidence of TK admixture (Lavue5, Lwa1-3, Blang) lack sacred pillars but engage in rituals honoring the spirit shrine and Buddhism [29]. This suggests that the introduction of Buddhism through contact with TK-speaking people has led to a decrease in spiritual practices among the latter group. Additionally, the degree of contact and admixture with TK people correlates with geographic locations, with populations in lowland river plains exhibiting a higher degree of TK admixture compared to those in mountainous areas (Fig. 1 and 3).

#### Blang

The Blang are another ethnic group that uses the Palaungic language and is sometimes classified as the “Lua” ethnicity. Their original homeland was in the area along the border between Myanmar and China. They are also known by other ethnonyms such as Bulang, Sam Tao, and Tai Loi. While there is no clear evidence of the Blang people’s migration to Thailand, it is estimated that they began moving there less than 100 years ago [8]. In the past 20-30 years, the Blang people have been identified as part of the “Lua” ethnic group by Thai government officers, being recognized as one of the 13 expanded official hill-tribes of Thailand, which includes the Akha, Dara-ang, Hmong, Htin, Iu-Mien, Kachin, Karen, Khamu, Lahu, Lisu, Lua, Mlabri, and Shan [27].

Using genome-wide markers, we have found that the genetic structure of the Blang people is similar to that of the Lwa group, mainly comprising genetic components similar to the others in AA family but with admixture from TK gene pool (Fig. 3). The Blang people are an ethnic group that has had close historical interactions with other ethnic groups, particularly the TK-speaking Shan (Tai Yai) ethnicity, mainly through the trade of various daily life items, which has likely led to assimilation between them.

## Conclusion

We utilized genome-wide markers to analyze the genetic clustering patterns of AA-speaking ethnic groups in northern Thailand. Our findings align with linguistic classifications, revealing distinct genetic profiles among the three branches of the Mon-Khmer subfamily of the AA family: Monic, Khmuic, and Palaungic. Despite confusion in using the ethnic name “Lua” to identify groups from both Khmuic and Palaungic branches, our genomic data clarifies that Khmuic-speaking Lua communities exhibit genetic differences from Palaungic-speaking Lavue and Lwa groups.

Within the Palaungic branch, the Dara-ang stand out as the most genetically distinct, retaining some remnants of ancient genetic composition. Lavue populations, mainly residing in mountainous areas, display a genetic makeup unique to the AA family and share genetic closeness with the Karenic of the ST family. On the other hand, the Lwa and Blang, residing in lowland river valleys, show genetic components resulting from admixture with TK-speaking ethnic groups. While self-identification ethnonyms and linguistic classifications can roughly categorize Palaungic-speaking populations in northern Thailand, we propose that genetic relationships are more closely linked to spiritual practices (the presence or absence of sacred pillars in communities) and geographic locations (mountainous or lowland areas).

## Materials and Methods

### Ethical Statement

The study received ethical approval from the Human Experimentation Committee of the Research Institute for Health Sciences at Chiang Mai University, Thailand (Ethical Clearance Certificate No. 45/2023). Throughout the research, we ensured the protection of participants’ rights and identities, adhering to all relevant guidelines and regulations outlined in the experimental protocol on human subjects as per the Declaration of Helsinki. Prior to interviews and sample collection, written informed consent was obtained from all volunteers.

### Sample collection and Quality Control

We collected samples from 95 unrelated individuals living in 5 villages in Chiang Mai and Mae Hong Son provinces of northern Thailand. The participants were healthy subjects over 20 years old, of Palaungic-speaking ethnicity, and had no known ancestors from other recognized ethnic groups for at least three generations. We collected personal data through form-based oral interviews for endonyms, self-reported unrelated lineages, linguistics, and migration histories. Buccal specimens were collected, and genomic DNA was extracted using the Gentra Puregene Buccal Cell Kit (Qiagen, Germany), following the manufacturer’s instructions. Detailed sample information is listed in Table 1, and the geographic locations of sampling are shown in Fig 1.

Note that, to reduce confusion regarding terminology and classification under the “Lua” ethnicity, we will primarily use the self-identifying endonyms that residents use to refer to themselves and will further refer to names used in previous reports.

### Genotyping and data preparation

Genotyping was carried out using the Affymetrix Axiom Genome-Wide Human Origins array [30] at AtlasBiolab in Germany. A total of 93 samples were genotyped for 629,721 genetic markers on the hg19 version of the human reference genome, with a genotype call rate of at least 97%. We used PLINK version 1.90b5.2 [31] to remove genetic markers and individuals with more than 5% missing data, as well as to exclude mitochondrial DNA and sex chromosomes from our analysis. Loci that did not pass the Hardy-Weinberg equilibrium test (p-value <0.00005) or had more than 5% missing data within any population were also excluded. Additionally, we used KING 2.3 [32] to assess individual relationships and removed one person from each pair of first-degree kinship. After these quality control steps, we had data from 89 Palaungic-speaking individuals with 617,132 genetic markers in total (Table 1).

We utilized the “bmerge” function in PLINK version 1.90b5.2 to combine our newly genotyped Palaungic-speaking data with genome-wide SNP data from modern and ancient populations in East and Southeast Asia, as well as reference populations like the Mbuti and French from the Allen Ancient DNA Resource (AADR) version 54.1 [33]. Additionally, we incorporated data obtained from previous studies [12,18,20,34–37]. We then assessed the data quality using PLINK version 1.90b5.2 and excluded SNPs with more than 5% missing data, less than 15,000 SNP positions, or those not in Hardy-Weinberg equilibrium with a significance level of p-value < 0.00005. This resulted in a dataset comprising 2,390 individuals from 263 populations (S1 and S2 Table) with 499,039 SNP positions available for further analysis.

### Population structure and relationship analyses

The genetic structure and relationships of the studied sample were analyzed by pruning SNPs in the same linkage disequilibrium with an r^2^ value greater than 0.4 within windows of 200 variants and a step size of 25 variants using the “indep-pairwise” command in PLINK version 1.90b5.2. After excluding the Mbuti and French populations used as outgroups, there were 219,244 SNP positions available for analysis from a total of 2,355 individuals. Principal Component Analysis (PCA) was carried out using smartpca, a part of the EIGENSOFT package [38]. We used all default parameters along with additional parameters of lsqproject: YES and autoshrink: YES. The ancient DNA data, which differs genetically from modern populations, was projected.

The ADMIXTURE, a model-based clustering algorithm for ancestry estimation method, was utilized to investigate the genetic composition of a merged dataset comprising 2,390 individuals from 263 modern and ancient populations. This analysis aimed to discern genetic structures and infer ancestral origins with respect to ethnicities and linguistic groups. We employed ADMIXTURE version 1.3.0 [13], varying the number of assumed ancestral components (K) from 2 to 12, and conducted 100 bootstrap iterations with different random seeds. The optimal K value was determined by assessing the lowest cross-validation error and the highest log-likelihood using 10-fold cross-validation. Data visualization was accomplished using PONG software version 1.4.7 [39] and R software [40].

Furthermore, we examined drift and excess ancestry sharing within the studied populations by calculating *f3* and *f4*-statistics using ADMIXTOOLS version 5.1 [30], in conjunction with admixr version 0.7.1 [41]. The Mbuti population was used as an outgroup for modern-modern population analysis, while the French population served as an outgroup for modern-ancient population analysis. The data were then visualized using the pheatmap package in R software version 3.6.0 to generate a heatmap.

### Genetic ancestry analyses

To investigate possible ancestral origins and genetic exchanges, we initially created a TreeMix-based tree involving our focus populations and several Asian ancestral groups. The African Mbuti, European French, and Indian populations were utilized as outgroups for this analysis. To streamline the tree structure, we organized the TK populations based on their linguistic subfamilies, whereas the ST, AN, and HM populations were grouped according to their geographic location (East or Southeast Asia). Using the TreeMix version 1.12 program [15], we reconstructed a phylogenetic tree with migration events ranging from 0 to 3 using 10 independent runs and then selected the topology with the highest likelihood.

We proceeded to create an admixture graph, selecting backbone populations from various language families in Southeast Asia as outlined in [20]. The Atayal, Dai, Cambodian, Miao, and Naxi were chosen as representative ethnic groups that speak AN, TK, AA, HM, and ST languages, respectively. We then incorporated our interest in AA-speaking and the specific case between the Lavue and Karen populations into the graph. For modeling the admixture graph, we utilized a dataset of 499,039 SNPs from modern populations as input for ADMIXTOOLS 2 [42]. Initially, we calculated pairwise *f2* statistics between groups using the “extract_f2()” function with specific parameters; “maxmiss = 0” (no missing SNPs for calculation), “useallsnp: NO” (allowing no missing data), and “blg = 0.05” (setting SNP block size at 0.05 morgans). Subsequently, we derived allele frequency products from the computed *f2* blocks using “f2_from_precomp()”.

Next, we explored the best-fitting admixture graph by running ten independent iterations of “find_graphs()” for each scenario. Among the 100 independent runs, we selected the one with the lowest score, calculated based on residuals between expected and observed f-statistics given the data. To validate the selected graph, we tested it using “qpgraph()” with parameters “numstart = 100, diag = 0.0001, return_fstats = TRUE”, checking if the worst-fitting Z score absolute value was below 3. Starting with no migrations (numadmix = 0), we incrementally added migrations until reaching a fitting graph, which we considered the best-fitting graph for that specific scenario.

## Data Availability

The datasets generated and analyzed during the current study are available in the Zenodo repository (https://doi.org/10.5281/zenodo.10957983)

## Acknowledgements

We would like to express our gratitude to all volunteers, related government officers, and abbot of Khun Khong Temple in Chiang Mai province, for their efforts in collecting information and samples. We are grateful to Dr. Pittayawat Pittayaporn and the students of Chulalongkorn University for their participation in observing our sample collection and laboratory processes for this study. Special thanks to Dr. Dang Liu for providing valuable advice on computational analyses. J.K. acknowledges Mentor Wiluck Sripasang for inspiring the initiation of this project. This research project was supported from the Fundamental Fund 2023, Chiang Mai University, Thailand (FF66/050). Additionally, W.K. and M.S. received funding from Naresuan University under the Global and Frontier Research University Fund (R2566C051).

## Author Contributions

J.K., W.K., M.S., A.I. conceived and designed the project; J.K., T.S., M.D. and S.S. collected samples and generated data; J.K., T.S., and W.K involved in data analyses; J.K. wrote the main manuscript with input from all co-authors.

## Competing interests

The authors declare no competing interest.

## Supplementary information

**S1 Fig. The cross-validation values of ADMIXTURE ranging from K=2 to 12.**

**S2 Fig. ADMIXTURE diagram showing the genetic components of various East and South Asian populations.** K values ranging from 2 to 10 divided into groups from K=2 to K=12 for modern populations (left) and ancient DNA (right). Each individual is represented by a bar divided into K colored segments, indicating their estimated membership fractions in each of the K ancestry component. Populations are separated by black lines and their names are labeled in colors showing different linguistic family.

**S3 Fig. Heatmap showing allele sharing profiles based on *f3* statistics comparing the AA speaking populations with other Asian ethnic groups**. The colored bar on the top-right indicates the statistical values, while that on the low-right indicates the linguistic family of each ethnic group.

**S4 Fig. Heatmap showing allele sharing profiles based on *f3* statistics comparing the AA speaking populations with ancient DNA.** The colored bar on the right indicates the statistical values.

**S5 Fig. The diagram illustrates the relationship between populations analyzed using *f4*-statistics, comparing modern populations with ancient DNA in Southeast Asia.** *f4* statistics comparing the AA speaking populations labeled on the left to the ancient samples from Southeast Asia on the top grey bar. Z-scores are for *f4* (ancient sample, Han Chinese; ethnic population, French). The vertical grey lines denote 0. Empty circles denote nonsignificant Z-scores (|Z|=<3) and solid circles denote significant Z-scores (|Z|>3)

**S6 Fig. The TreeMix diagram for the AA-speaking ethnic group in northern Thailand and other modern populations in East and Southeast Asia.** (A) without migration event (B) one migration event (C) two migration event (D) three migration event. Populations are labeled with different colors based on their language family.

**S1 Table General information concerning populations and samples studied.**

**S2 Table Summarized data of populations using for comparison.**

**S3 Table The result of the *f3* (X, Y: Outgroup), where Outgroup = Mbuti**

**S4 Table The result of the *f4* (Dara-ang, Palaungic-speaking group: Asian, Outgroup), where Outgroup = Mbuti**

**S5 Table The result of the *f4* (ancient samples, Han Chinese: ethnic groups, Outgroup) with transversion only**

## Notes

### Competing Interest Statement

The authors have declared no competing interest.

